# Analysis of variance when both input and output sets are high-dimensional

**DOI:** 10.1101/2020.02.15.950949

**Authors:** Gustavo de los Campos, Torsten Pook, Agustin Gonzalez-Raymundez, Henner Simianer, George Mias, Ana I. Vazquez

## Abstract

**Motivation:** Modern genomic data sets often involve multiple data-layers (e.g., DNA-sequence, gene expression), each of which itself can be high-dimensional. The biological processes underlying these data-layers can lead to intricate multivariate association patterns.

**Results:** We propose and evaluate two methods for analysis variance when both input and output sets are high-dimensional. Our approach uses random effects models to estimate the proportion of variance of vectors in the linear span of the output set that can be explained by regression on the input set. We consider a method based on orthogonal basis (Eigen-ANOVA) and one that uses random vectors (Monte Carlo ANOVA, MC-ANOVA) in the linear span of the output set. We used simulations to assess the bias and variance of each of the methods, and to compare it with that of the Partial Least Squares (PLS)–an approach commonly used in multivariate-high-dimensional regressions. The MC-ANOVA method gave nearly unbiased estimates in all the simulation scenarios considered. Estimates produced by Eigen-ANOVA and PLS had noticeable biases. Finally, we demonstrate insight that can be obtained with the of MC-ANOVA and Eigen-ANOVA by applying these two methods to the study of multi-locus linkage disequilibrium in chicken genomes and to the assessment of inter-dependencies between gene expression, methylation and copy-number-variants in data from breast cancer tumors.

**Availability:** The Supplementary data includes an R-implementation of each of the proposed methods as well as the scripts used in simulations and in the real-data analyses.

**Supplementary information:** Supplementary data are available at *Bioinformatics* online.

## 1 Introduction

Modern genomic data sets often combine information from multiple data-layers, each of which itself can be high-dimensional. Examples of this include data sets comprising of information from several omics, or data sets combining genomic information with high-throughput phenotyping (e.g., crop-imaging imaging, milk infrared spectra data). The biological processes underlying each of the data-layers can induce complex dependencies between features within (e.g., linkage disequilibrium among single nucleotide polymorphisms, SNPs) as well as between layers (e.g., association between DNA and gene expression, GE). The main goal of this study is to develop and to evaluate methods for quantifying multivariate-associations in settings in which both the input and output sets are high dimensional.

The methods proposed in this study can be used to answer ubiquitous questions such as: How much of the inter-individual differences in whole-genome sequence genotypes can be predicted using a low-density SNP array? What proportion of variance in GE can be explained by differences in DNA methylation (ME)? How much of the variance in image-phenotypes can be predicted from DNA genotypes?

Canonical Correlation Analysis (CCA, Mardia, T., and Bibby, 1979), Multivariate-Analysis of Variance (MANOVA, Rencher and Christensen, 2012), and Reduced Rank-Regressions, e.g., Partial Least Squares (PLS, Wold, Sjöström, and Eriksson, 2001) are three methodologies often used to assess associations in multi-dimensional problems. However, these approaches have limitations that make some of them inadequate for estimating the proportion of variance explained when both the output and input layers are high-dimensional.

**Canonical Correlation Analysis** extends the concept of the correlation between two random variables to cases involving two multidimensional data sets; however, correlation is symmetric by nature. Therefore, CCA cannot address questions regarding proportion of variance explained when the proportion of variance of one set (e.g., ***X***) that is explained by another set (***W***) is not equal to the reciprocal (i.e., proportion of variance of ***W*** that can be explained by ***X***). In many multi-layered data sets we do not expect to have a symmetric variance-decomposition (we illustrate this below using simulations and experimental data).

**Multivariate Analyses of Variance** (MANOVA, e.g., Krzanowski and J. 1988) is another approach that can be considered. However, MANOVA is based on least-squares projections; therefore, the methodology is not well-suited for cases when data is high dimensional, including rank-deficient cases. Most of the problems we are focusing on involve high-dimensional data where the number of features exceeds sample size, thus, making least-squares methods such as MANOVA inadequate.

**Reduced-rank regressions** (e.g., Izenman, 1975) **and penalized multivariate analysis methods** (e.g., Witten, Tibshirani, and Hastie 2009) are often used to analyze high-dimensional data. However, the results that one can obtain using regularized methods (including reduced-rank and penalized methods) rely on regularization decisions (e.g., the number of dimensions used in PLS or CCA, or the sparsity parameters in sparse CCA) which cannot be made using the likelihood. Thus, these parameters are often tuned to maximize prediction accuracy in testing sets. This approach is not necessarily optimal for inferences.

Therefore, to overcome the limitations of existing methods, we developed two approaches for estimation of proportion of variance explained when both the input and output sets are high-dimensional. Our approaches use random effects models to estimate the proportion of variance of (independent) vectors in the linear span of an output layer that can be explained by regression on an input layer. We considered two methods for generating a sequence of independent vectors in the linear span of the output layer: A Monte Carlo method (MC-ANOVA) which uses random vectors, and one based on eigenvectors (Eigen-ANOVA).

## 2 Methods

Let ***X*** and ***W*** denote two matrices holding data for *n* individuals (rows) and *p* (***X***) and *q* (***W***) features in columns. For instance, ***X*** may be a matrix with genotypes codes at *p* SNPs and ***W*** may be a matrix providing GE levels assessed at *q* genes.

The columns of ***X*** = {***x***_1_, ***x***_2_, …, ***x***_*p*_} and of ***W*** = {***w***_**1**_, ***w***_2_, …, ***w***_*q*_} can be viewed as axes spanning two linear spaces (*L*_*X*_ and *L*_*W*_, respectively). The linear spans of ***X*** and ***W*** consist of all the vectors that can be obtained by forming linear combinations of the columns of each of these sets, that is 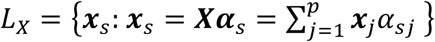 and 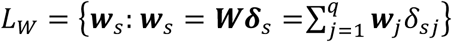, for all real-valued vectors ***α***_*s*_ = {*α*_*s*1_, …, *α*_*sp*_} and ***δ***_*s*_ = {*δ*_*s*1_, …, *δ*_*sp*_}.

For each vector ***x***_*s*_ ∈ *L*_*X*_, one can estimate the proportion of variance that can be explained by linear regression on *L*_*W*_ using a model of the form

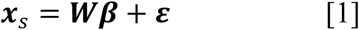

For cases where *q* is large, the proportion of variance of ***x***_*s*_ that can be explained by regression on *L*_*X*_ 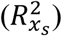 can be estimated by regarding both ***β*** and ***ε*** as Gaussian independent random variables, 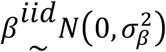 and, 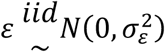. Upon appropriate scaling of the columns of 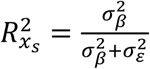 can be interpreted as the proportion of variance of the phenotype that could be explained by regression on the features included in ***X***. The variance parameters involved (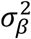 and 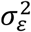) can be estimated using Bayesian or Likelihood methods (e.g., restricted maximum likelihood, REML, Patterson and Thompson 1971).

In the preceding paragraph we describe how one can estimate the proportion of variance of a vector in *L*_*X*_ (***x***_*s*_) that can be explained by regression on the linear span of ***W***. Our goal is to generalize this to all vectors in *L*_*X*_. However, *L*_*X*_ contains an infinite number of vectors; therefore, some approximation is needed.

Perhaps the most natural approach for estimating the proportion of variance of vectors in *L*_*X*_ that can be explained by regression in *L*_*W*_ consists of regressing each of the columns of ***X*** on ***W***. Such an analysis would produce a sequence of R^2^ estimates 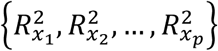, and the average R^2^, 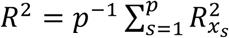, could be used to estimate the overall proportion of variance of ***X*** that could be explained by regression on ***W***. However, one limitation of this approach is that the columns of ***X*** are often not independent. Many features may cluster (e.g., genes may be co-expressed, or SNPs may be in high linkage-disequilibrium) leading to groups of highly unbalanced sizes. When some features are highly-correlated, the simple average of individual R^2^-values may lead to inaccurate (even biased) estimates. Furthermore, when ***X*** is ultra-high dimensional (e.g., hundreds of thousands or million features) estimating *R*^2^, on-feature-at-a-time will be computationally challenging. To address these problems, we discuss two methods that use independent vectors in the span of the output set.

### Method 1: Monte Carlo Analysis of Variance (MC-ANOVA)

Since *L*_*X*_ is infinite, one we cannot exhaustively estimate 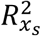 for all vectors in *L*_*X*_. However, we can ‘explore’ the linear span of the output set by generating random vectors in *L*_*X*_ of the form ***x***_1_ = ***Xα***_*s*_, where ***α***_*s*_ is sampled from some distribution. This can be repeated for a large number of vectors in *L*_*X*_ to produce a sequence of estimates 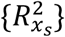, and the resulting sequence can be used to estimate the average proportion of variance explained as well as other features of the distribution of the sequence. The method is summarized in Box 1. Importantly, if ***α***_*s*_ and ***α***_*s*′_ are independent, so will be ***x***_*s*_ and ***x***_*s*′_.

##### Box 1. Monte Carlo Analysis of Variance (MC-ANOVA)

1. Draw a random vector ***α***_*s*_ from a proper multivariate distribution
2. Form the linear combination ***x***_*s*_ = ***Xα***_*s*_
3. Estimate the proportion of variance of 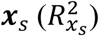 using a random effects model (expression [1]) with variance parameters estimated using either Bayesian or likelihood-type methods.
4. Repeat 1-3 for a large number of times.
5. Obtain a global R^2^ estimate by averaging the 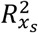’s in the sequence.

In Box 1 we did not specify how the ***α***_*s*_ are generated. One possibility is to sample these coefficients from distributions with support in *R*^*p*^ (e.g., *p*-variate Gaussian). Alternatively, one could sample sparse vectors of coefficients from finite mixtures with a point of mass at zero. The possibility of using different process for generating random vectors in *L*_*X*_ gives the MC-ANOVA a great deal of flexibility–we will further explore that flexibility in greater detail in the case studies presented further below.

### Method 2: Regression using orthogonal basis (Eigen-ANOVA)

An orthogonal basis for the row-space of ***X*** can be obtained from the singular-value decomposition of ***X*** = ***U***_*X*_***D***_*X*_***V***′_*X*_, where ***U***_*X*_ and ***V***_*X*_ are the left- and right-singular vectors of ***X*** respectively, and ***D***_*X*_ is a diagonal matrix with the singular values of ***X*** in the diagonal. Both ***U***_*X*_ and ***V***_*X*_ are orthonormal, thus ***U***′_*X*_***U***_*X*_ = ***I*** and ***V***′_*X*_***V***_*X*_ = ***I***. Each vector in *L*_*X*_ can be represented as a linear combination of the left-singular vectors of ***X***. Therefore, our second method estimates the proportion of variance of each of the left-singular vectors of ***X*** that can be explained by regression on ***W***, and produces a global R^2^ estimate using a weighted sum of the R^2^ estimated for each singular vector (Box 2).

##### Box 2. Eigen-ANOVA

1. Generate an orthogonal basis for *L*_*X*_; for instance, compute the singular-value decomposition of ***X*** = ***U***_*X*_***D***_*X*_***V***′_*X*_ where ***U***′_*X*_***U***_*X*_ = ***I*** and ***V***′_*X*_***V***_*X*_ = ***I*** are orthonormal basis for the row- and column-space of ***X*** respectively, and ***D***_*X*_ = *Diag*{*d*_*i*_} is a diagonal matrix with the singular values of ***X*** in its diagonal.
2. Regress each of the left-singular vectors on *L*_*W*_ using a linear model such as that in expression [1] with ***u***_*i*_ = ***x***_*s*_, and estimate the proportion of variance of each vector that can be explained by regression on *L*_*W*_, 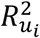.
3. Estimate the global proportion of variance of vectors in *L*_*X*_ that can be explained by regression on *L*_*W*_ using 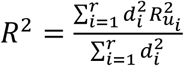.

## 3 Results

In this section we first present results from simulations designed to detect bias on estimates derived from each of the methods, and to compare the bias of the proposed methods with that of the PLS–a method commonly used in multivariate-high-dimensional regressions.

### 3.1 Statistical properties assessed via simulations

**Data** were simulated using genotypes from a wheat data set generated by the International maize and Wheat Improvement Center (CIMMYT). Briefly, the data set provides genotypes at 1,279 molecular markers assessed in 599 wheat inbred lines. Further details about this data set are provided by de los Campos et al. (2009). The data set is available with the BGLR R-package (de los Campos and Perez-Rodriguez, 2015). The scripts used to conduct these simulations are provided in the Supplementary Data (File S1). The REML estimation algorithm used to implement the MC- and Eigen-ANOVA is also provided in File S1. (We also tested the estimator proposed by Schreck et al. 2019; the results were very similar to the REML estimates presented below.) As benchmark, we used the PLS regression method. Briefly, we regressed the output matrix (***X***) on the input data (***W***) using the plsr function of the pls R-package (Mevik, Wehrens, and Liland 2019). The number of orthogonal basis used was determined using cross-validation–we choose the number of components that maximized prediction accuracy. The R-squared was then computed for each feature using the entire data set, and an overall R-squared was obtained by averaging the R-squared obtained for each of the columns of ***X***. The implementation can be found in File S1.

#### Simulation 1

In our first simulation we use the genotypes of the wheat data set as the input set: ***W*** = [***w***_1_, …, ***w***_*q*_]. The ‘output’ set, ***X*** = [***x***_1_, …, ***x***_*p*_], was simulated using: ***x***_*i*_ = ***w***_*i*_ + ***δ***_*i*_ where *i=1,…,1,279* indexes molecular markers and ***δ***_*i*_ is an *n*-dimensional vector of *iid* (independent and identically distributed) random draws from a normal distribution with zero mean and a variance parameter, such that the proportion of variance of ***x***_*i*_ explained by ***w***_*i*_ was, 0, 0.1, 0.3, 0.5, 0.8, 0.9 and 1. Here, 0 represents the case of complete independence (***x***_*i*_ = ***δ***_*i*_) and 1 the case of perfect concordance (**X**=**W**).

We conducted a total of 1,000 Monte Carlo (MC) replicates per scenario. The input set (***W***) did not change across MC samples, however, the output set, ***X***, varied across MC replicates due to the noise term (***δ***_*i*_). For each simulated data set we then estimated the proportion of variance of **X** explained by regression of ***W*** using random effects models, with variance parameters estimated using REML (Patterson and Thompson 1971).

The Monte Carlo method estimated the proportion of variance of ***X*** explained by ***W*** without any noticeable bias (**Table 1**). However, the regression of the eigenvectors of ***X*** on ***W*** produced estimates that, in some scenarios, were downwardly biased. The presence of bias was evident in scenarios where the true proportion of variance of ***X*** explained by ***W*** was greater than 0.5. Further inspection of the results for individual MC replicates suggested that the bias of the Eigen-ANOVA method was likely due to a relatively large number of ‘corner’ solutions (zero estimated proportion of variance) which were common for high-order eigenvectors (i.e., those with small eigenvalue)–we illustrate this in an analysis of multi-omic cancer data further below. The R^2^ estimates obtained with PLS also had noticeable biases which, in most scenarios, were larger in absolute value than the bias estimates obtained with Eigen-ANOVA.

**Table 1.** Average (SD) estimate of the proportion of variance explained by simulation scenario (first column) and estimation method (Simulation 1).

| Simulated Proportion<br>of Variance Explained | Monte Carlo-ANOVA | Eigen-ANOVA | PLS |
| --- | --- | --- | --- |
| 0.0 | 0.0082<br>(0.0028) | 0.0081<br>(0.0006) | 0.0017<br>(0.0001) |
| 0.1 | 0.1002<br>(0.0083) | 0.0983<br>(0.0019) | 0.0478<br>(0.0034) |
| 0.3 | 0.2991<br>(0.0108) | 0.3020<br>(0.0028) | 0.2412<br>(0.0075) |
| 0.5 | 0.4992<br>(0.0102) | 0.5054<br>(0.0028) | 0.4865<br>(0.0076) |
| 0.8 | 0.8006<br>(0.0055) | 0.7857<br>(0.0017) | 0.8451<br>(0.0036) |
| 0.9 | 0.9012<br>(0.0033) | 0.8685<br>(0.0011) | 0.9403<br>(0.0016) |
| 1.0 | 1.0000<br>($<.0001$) | 0.9377<br>($<.0001$) | 0.9988<br>($<.0001$) |

#### Simulation 2

We designed a second simulation to consider the case where one of the sets (***X***) was included in the other set (***W***). We achieved this as follows:

- We set ***X*** to be a matrix containing a subset of the wheat genotypes. Specifically, we sampled at random 128 (10%), 256 (20%), 640 (50%), 895 (70%) or 1151 (90%) of the available diversity arrays technology (DArT) markers and formed with those genotypes our ***X*** matrix.
- Subsequently, ***W***=[***X***,***Z***], was formed by combining in a single matrix the columns of ***X*** and a matrix (***Z***) whose entries were filled with *iid* draws from standard normal distributions. The columns of ***X*** and ***W*** were all centered and scaled to unit variance; therefore, the proportion of variance of ***W*** that can be explained by ***X*** equals the ratio of the number of columns of ***W*** that are shared with ***X***. On the other hand, since *L*_*X*_ ∈ *L*_*W*_, the true proportion of variance of ***X*** explained by ***W*** was always 1.

In our second simulation study the MC-ANOVA method rendered nearly unbiased estimates of the proportion of variance of one set explained by the other (**Table 2**). However, the Eigen-ANOVA method and the PLS produced noticeable biases. In most cases, both methods were downwardly biased; however, the PLS had an upward bias in a few scenarios.

**Table 2.**
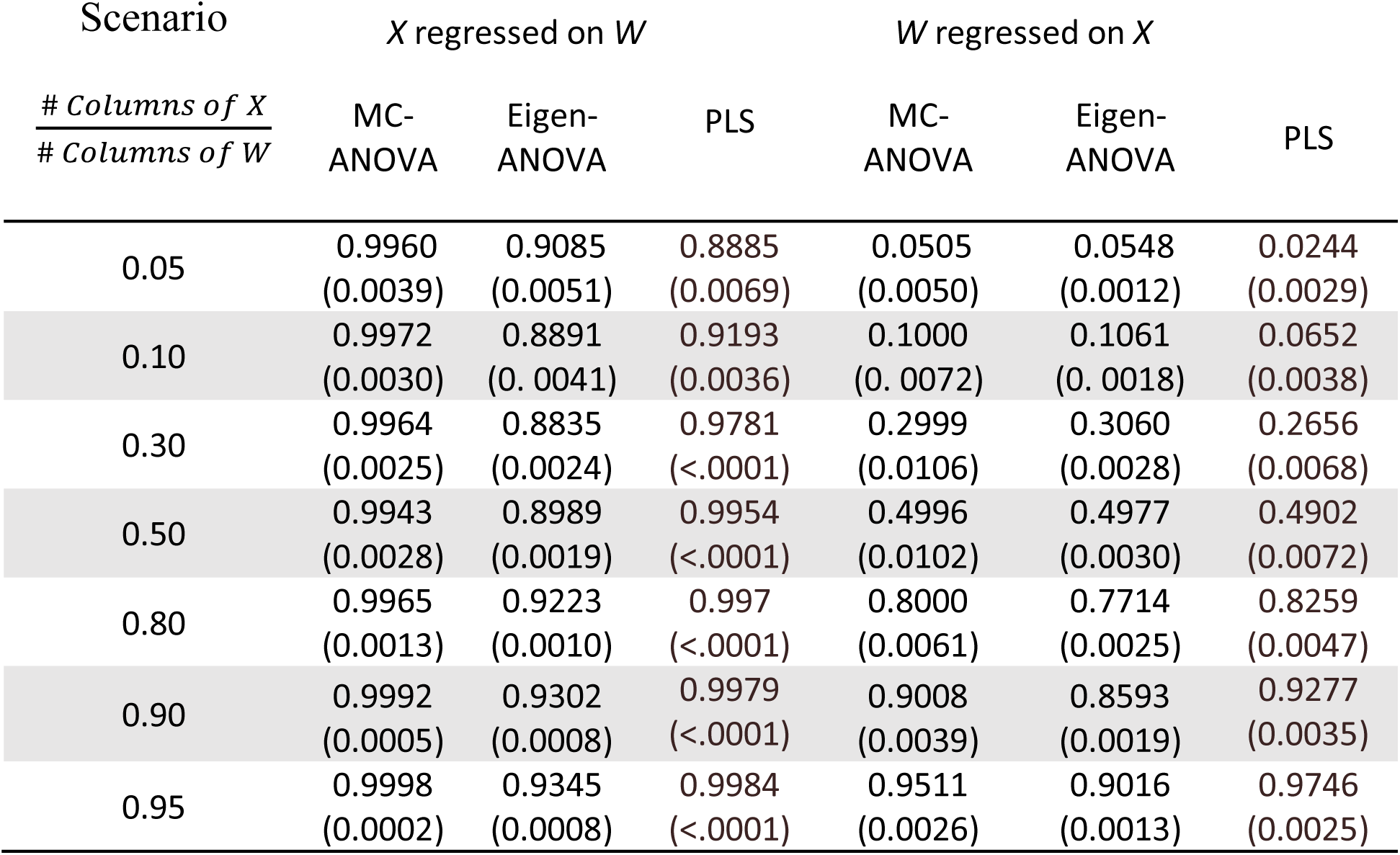
Average (SD) REML estimates of the proportion of variance explained by simulation scenario (first column) and estimation method (second simulation).

### 3.2 Applications to experimental data

We used the MC-ANOVA and Eigen-ANOVA to quantify the proportion of variance explained in two experimental data sets. The first data set contains a set of ultra-high-density (UHD, 1million+SNPs) SNPs derived from a combination of whole-genome sequencing (WGS) and imputation, and a set of (in-silico created) low-density SNP panels. We use this data set to assess the proportion of variance of UHD genotypes that can be captured and predicted using low-density SNP sets. The second data set involved three omic-layers (GE, ME, and copy-number-variants, CNV) of female breast cancer tumors. We used this data set to assess the proportion of variance at one omic that can be explained by another omic.

#### Case-study 1: Multi-locus linkage disequilibrium between ultra-high-density SNP genotypes and low-density SNP sets in chicken genomes

The continued reduction of genotyping and sequencing costs has led to a sustained increase in the number of loci that can be genotyped. In plant and animal breeding four typical genotyping options include: (i) customized low density arrays with anywhere between hundreds or a few thousand SNPs, (ii) commercial arrays of common SNPs with circa 50K (K=1000) SNPs, (iii) high density SNP arrays with a number of markers of ∼0.5M (M=million) SNPs and (iv) whole-genome sequence-derived SNP genotypes. The number of SNPs that can be derived from WGS varies between populations, but is usually of the order of tens of millions (UHD SNP genotypes). In recent years, several projects have produced large volumes of fully sequenced genomes for various agricultural species and model organisms. However, generating, storing, and fitting models with UHD-genotypes can be logistically, economically, and computationally challenging. Moreover, empirical evidence seems to suggests that using UHD SNP-genotypes does not lead to substantial gains in prediction accuracy relative to models trained using tens of thousands of SNPs (Erbe et al. 2013; Ober et al. 2012). This often leads researchers and the industry wondering: *How many SNPs are needed to capture (almost all) the information contained in UHD SNP genotypes?* We used the MC- and Eigen-ANOVA methods to address precisely this question.

***Data*** consisted of UHD SNP genotypes of 892 female and male chickens from six generations of a purebred commercial brown layer line of Lohmann Tierzucht GmbH. The genomes of 25 layers were sequenced at 8x read-depth, from the sequence data 4.92M (M=million) SNPs were derived. The remaining layers (n=867) were genotyped using the Affymetrix Axiom Chicken Genotyping Array (Kranis et al. 2013) which contains ∼600K (580,961) SNPs, and 336,224 passing quality control and filtering. The genotypes of these layers were then combined with those derived from WGS (4.92M SNPs) by first imputing the HD-data from the chip using BEAGLE 3.3.2 (Browning and Browning 2009) and afterwards imputing to UHD (4.92M SNPs) via MiniMac3 (Howie et al. 2012) with pre-phasing performed in BEAGLE 4.0. For details on the imputing procedure we refer to Ni et al. (2015). From the 4.92M SNPs we remove those with minor-allele-frequency smaller than 0.005 (0.5%) and pruned SNPs to guarantee a maximum R^2^ between adjacent SNPs of 0.99; these editing criteria left a total of 1.79M SNPs which were used for the subsequent analysis.

##### Proportion of variance at ultra-high-density genotypes explained by low-density SNP-sets

In a first analysis, the output space was the linear space (*L*_*X*_) spanned by the UHD SNP genotypes. The input set, (*L*_*W*_), consisted of low-density genotypes obtained by selecting *p* (*p*=500, 1K, 2K, 3K, 5K and 10K) evenly-spaced (in variant counts) SNPs. We estimated the proportion of variance captured by low-density panels using the Eigen- and MC-ANOVA. For the MC methods we sampled weights from *iid* standard normal distribution, 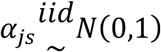 and then form a random vector in *L*_*X*_ using ***x***_*s*_ = ***Xα***_*s*_, where **X** is the matrix of UHD SNP-genotypes. These random vectors were then regressed on the lower-density SNP sets, and the proportion of variance explained was estimated using REML. This was repeated 1,000 times to estimate the distribution of proportion of variance of vectors in *L*_*X*_ explained by each of the low-density SNP-sets. For the Eigen-ANOVA we regressed each of the eigenvectors of the UHD SNP genotypes on the low-density panels.

According to the MC-ANOVA method, the panel containing 500 evenly-spaced SNPs captured about two thirds of the variance spanned by the UHD SNP genotypes (**Figure 1**). The proportion of variance of the UHD SNPs explained by low-density panels increased with the number of SNPs in the low-density panels reaching 100% with p=10K SNPs. The variance in the proportion of variance captured by low-density panels also decreased with the number of SNPs in the array (Figure 1). The Eigen-ANOVA yielded a very similar estimate of proportion of variance explained as the MC-ANOVA for *p*=500. However, for SNP-panels with more than 500 SNPs, the estimated proportion of variance obtained with the Eigen-ANOVA was systematically lower than the one obtained with MC-ANOVA. This agrees with what we found in the simulations where for high R-sq. the Eigen-ANOVA method gave downwardly biased estimates. (Note that while the MC-ANOVA yields both a point estimate and measures of dispersion (across random vectors) of R^2^, the Eigen-ANOVA only yields the point-estimates which are shown in Figure 1.)

**Figure 1.**
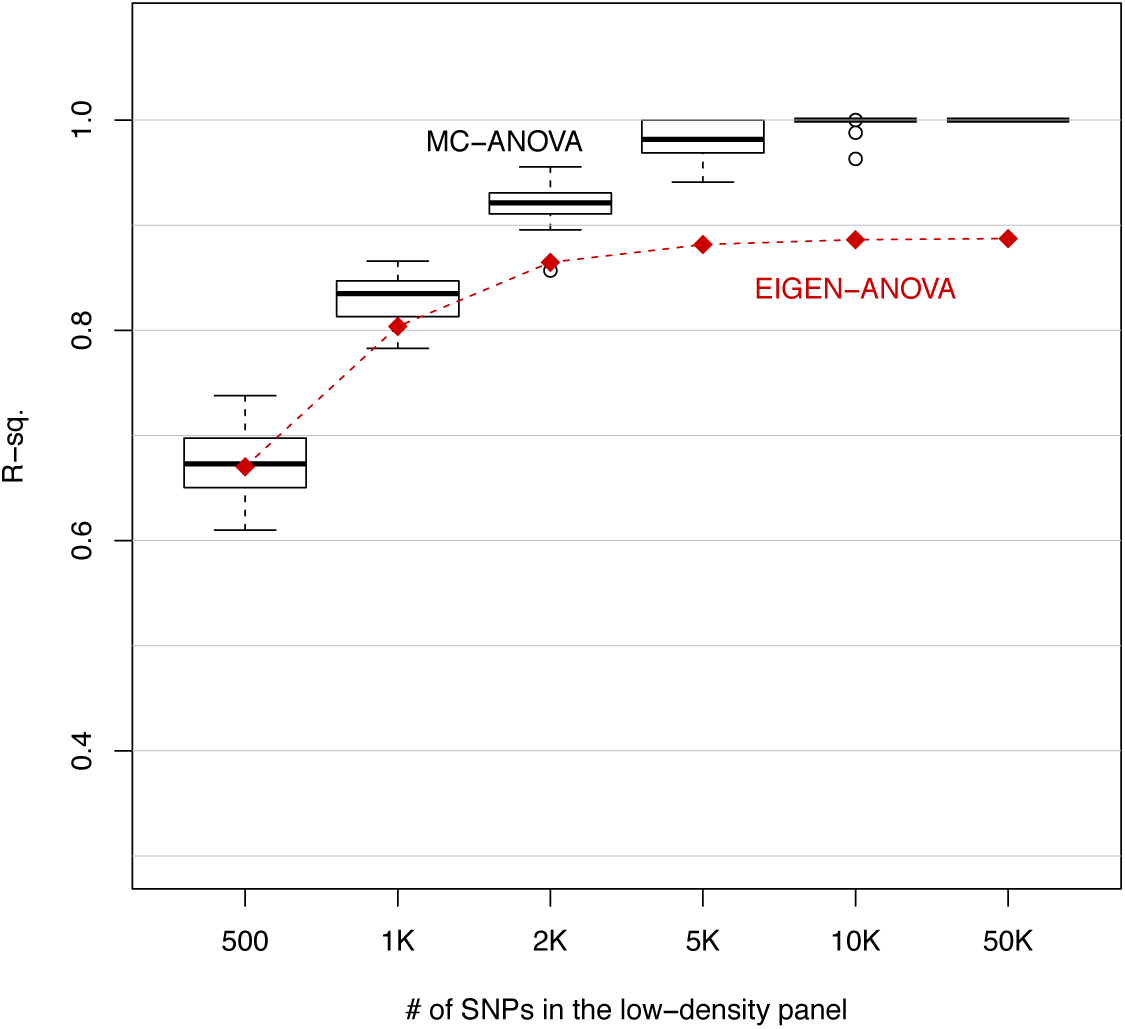
Proportion of the variance of whole-genome-sequence-derived SNPs (1.79 million) explained by SNP-panels consisting of 500, to 50K (K=1000) evenly-spaced SNPs.

##### Quantifying the effect of trait-complexity

In the previous application in the MC methods we drew random effect vectors that had weights (drawn from a normal distribution) on all the SNPs of the UHD set. However, for any trait, the vast majority of variants in the genome are expected to have no effect. The number of variants affecting any trait could vary from very few (simple traits) to hundreds or thousands (complex traits). Therefore, to explore the effect of the trait architecture on the distribution of the proportion of genetic variance of those traits that could be captured by low-density SNP panels, we repeated the previous analyses using random vectors that had *q (with q=5,10,50,500*) non-zero weights – the set of SNPs with non-zero weight were randomly sampled from the UHD-genotypes, and the weights of those SNPs were *iid* normal.

The estimated proportion of variance explained by regression on lower-density SNP panels were, on average, the same across “trait-architectures” (Figure 2). However, the dispersion about the estimated means was, as expected, much larger for simple traits. For “complex traits” with 500 “causal variants” the proportion of variance explained by regression on 10K or more SNPs was greater than 95% for all MC replicates. However, for simpler traits (e.g., 5 ‘causal variants’) we had some random vectors with proportion of variance explained smaller than 0.8.

**Figure 2.**
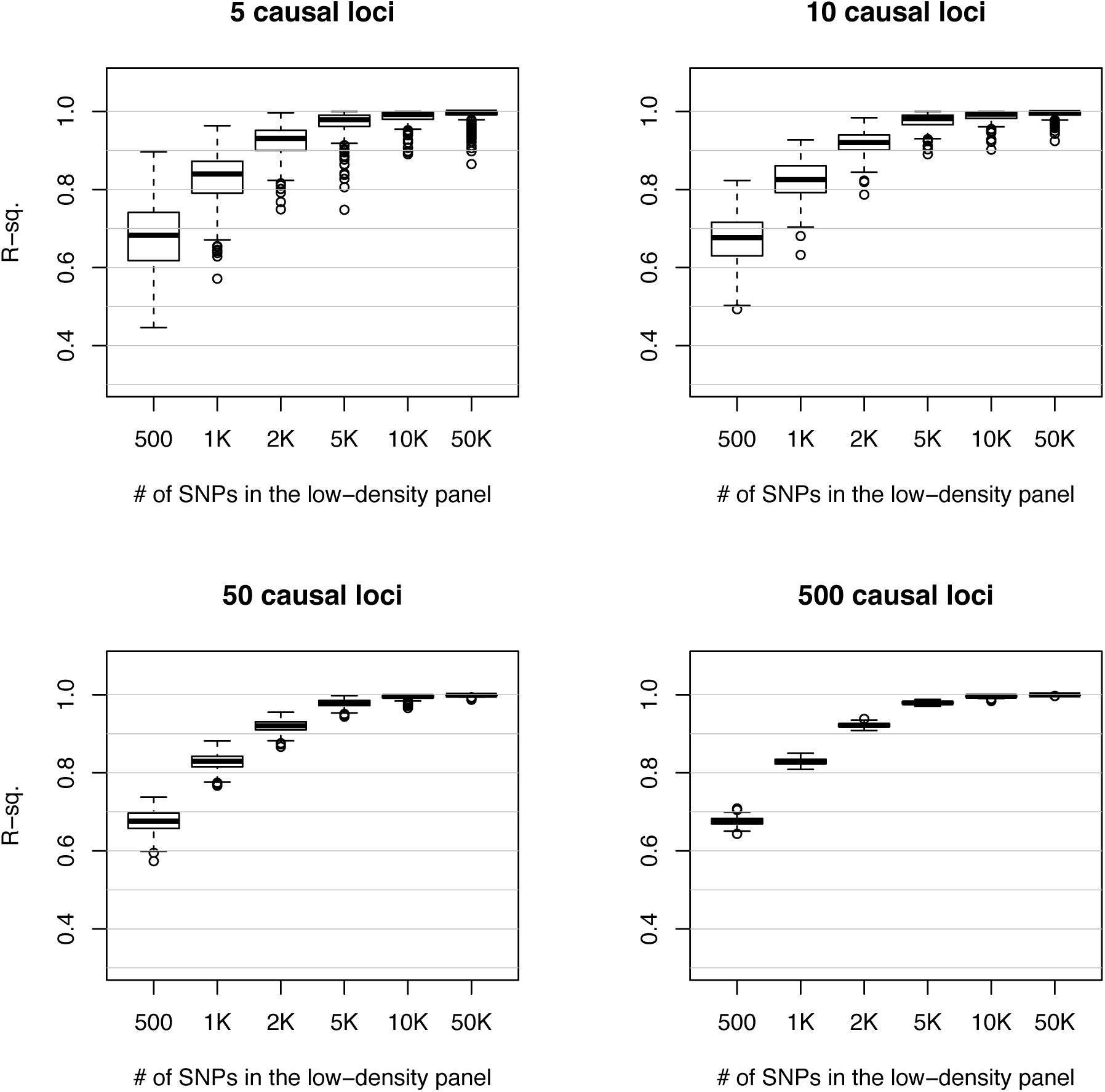
Proportion of variance of random vectors derived from ultra-high-density SNPs explained by regression on low-density SNP-panels, by number of loci used to form “genetic traits”.

#### Case Study 2: Proportion of variance explained in multi-omic data from breast cancer tumors

Cancerous processes involve the deregulation of signaling pathways controlling cell fate and progression, arising from the accumulation of genomic and epigenomics alterations across multiple genes (Vogelstein et al. 2013; Witte, Plass, and Gerhauser 2014). Genetic and epigenetic modifications can lead to changes in GE, which in turn can lead to changes in downstream (e.g., protein expression) and upstream (e.g., DNA, ME) processes, thus resulting in complex multivariate association patterns between multiple omic-layers.

##### Data

We used GE, ME and CNV data from breast cancer tumors to study multivariate associations between those three omics. Data (n=593) was from The Cancer Genome Atlas (TCGA), and consisted of samples from frozen primary breast cancer tumors from female patients.

Gene expression data (RNA-Sequencing counts) were determined using the Illumina HiSeq RNA V2 platform and DNA methylation profiles were determined using the Illumina HM450 platform. RNA-sequencing data were transformed using the natural logarithm and individual CpG site β-values were summarized at the CpG island level, using the maximum connectivity approach implemented in the WGCNA R package (Langfelder and Horvath 2008). The CpG island summaries were transformed into M-values (M=β/(1-β); Du et al. 2010) CNV profiles corresponded to gene-level copy number intensity derived from Affymetrix SNP Array 6.0 platform, using hg19 as reference.

##### Data editions

From each of the three omics we removed features with coefficient of variation smaller than 1% and those with proportion of missing values greater than 20%. The missing values that remained were imputed using a k-nearest neighbors clustering algorithm, with k = 3 (Hastie et al. 2016). After imputation, each feature was adjusted by batch effects using ComBat (Lazar et al. 2013). After applying the steps described above, the data set used in the analyses consisted of the (log-transformed) expression of 20,319 genes, 11,552 CVN-sites and ME intensity at 28,241 ME CpG islands.

##### Results

We used the MC- and the Eigen-ANOVA methods to estimate the proportion of variance of one omic that can be explained by regression on another omic; we did this for all pairwise omics combinations (GE∼ME, GE∼CNV, ME∼GE, ME∼CVN, CNV∼GE and CVN∼ME). Our results with the MC-ANOVA method indicate that the CNV data were completely explained by both GE and ME (Table 3). About 70% of the variance spanned by ME was explained by GE and vice versa. Finally, CNV explained a relatively small fraction of the variance spanned by either GE or ME. These results suggest that most CNVs have effects in both ME and GE and therefore, variation in CNV can be predicted by ME and GE. However, although there is association between CNV and both ME and GE, many other factors (e.g., environmental effects) seem intervene, thus, making the proportion of GE or ME explained by CNV relatively small (∼20%). Overall the MC- and Eigen-ANOVA methods yielded similar results. However, in cases involving high R^2^ (CNV∼ME, CNV∼GE, GE∼ME and ME∼GE) the Eigen-ANOVA method gave R^2^ estimates that were lower than those of the MC method. This pattern is consistent with what we observed in the simulation and in the analyses of chicken genomes.

**Table 3.**
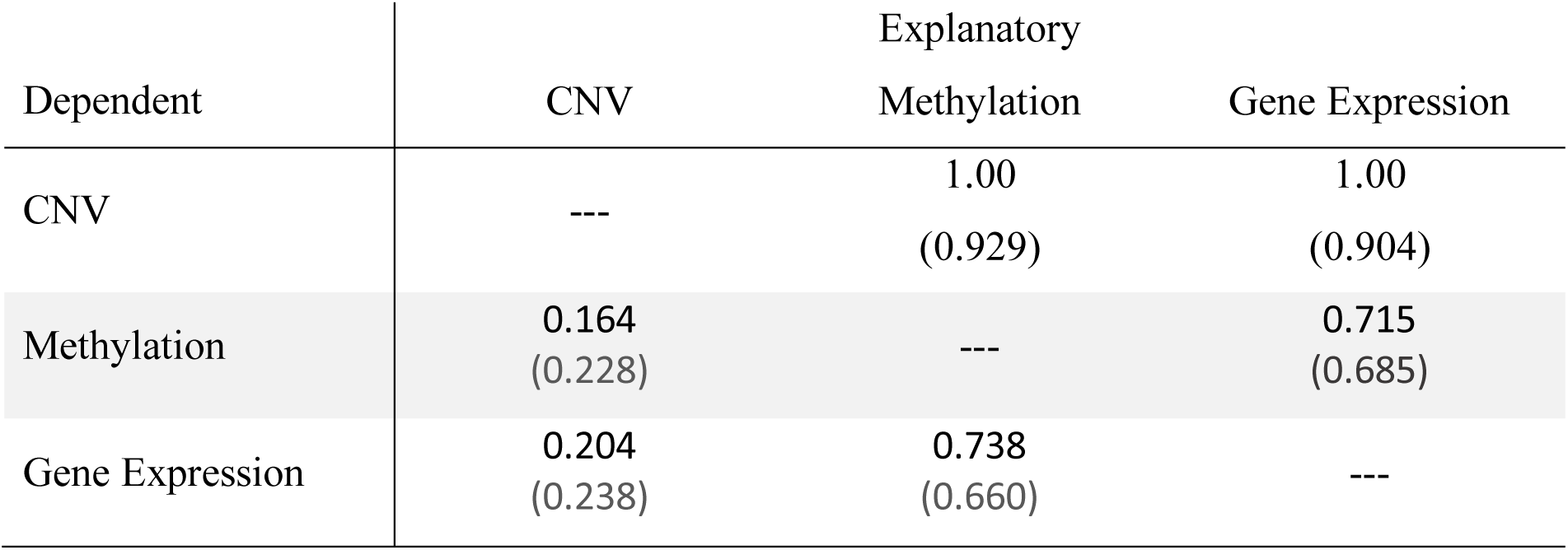
Proportion of variance of one omic explained by regression of the omic in each row on the omic in each column obtained with MC-ANOVA (Eigen-ANOVA).

| Dependent | Explanatory |  |  |
| --- | --- | --- | --- |
|  | CNV | Methylation | Gene Expression |
| CNV | --- | 1.00<br>(0.929) | 1.00<br>(0.904) |
| Methylation | 0.164<br>(0.228) | --- | 0.715<br>(0.685) |
| Gene Expression | 0.204<br>(0.238) | 0.738<br>(0.660) | --- |

Eigen-vector-specific R^2^ values obtained with the Eigen-ANOVA method (Figure 3) showed that the R^2^ values were, in most cases (except GE∼CNV and M∼CNV) very high (and in many cases very close to one) for the top-eigenvectors (i.e., those with high eigenvalue), and very small for eigenvectors associated with low eigenvalues. The transition in the R^2^ profile of individual eigenvectors showed a relatively sharp phase transition from R^2^ values near one to near-zero values. Overall, our results suggest a relatively good agreement in the patterns captured by the top-eigenvectors across omics.

**Figure 3.**
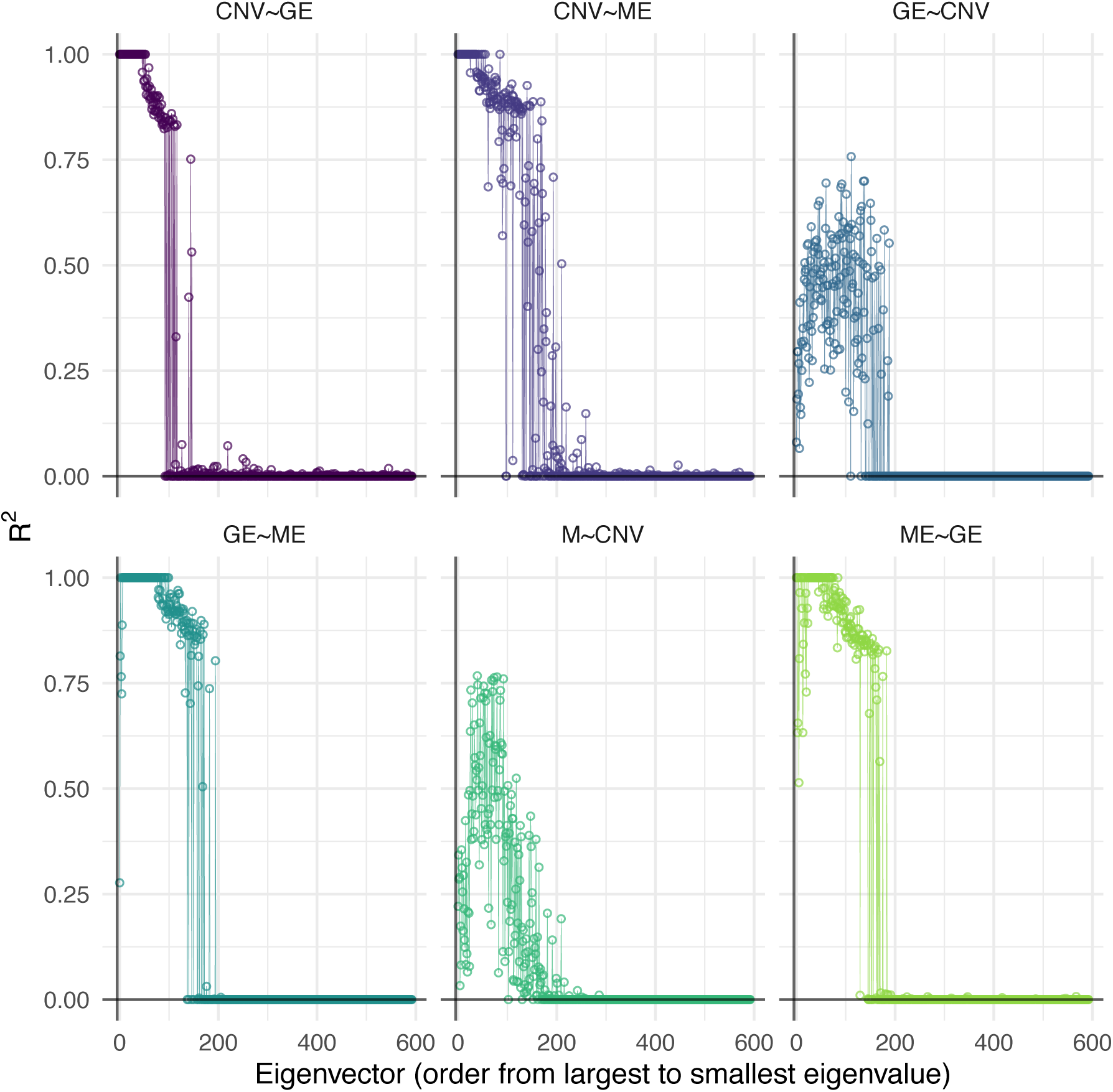
Proportion of variance of omic-derived eigenvectors of an omic-set explained by regression on a different omic-set. Points give the proportion of variance for individual eigenvectors. GE=Gene Expression, ME=Methylation, CNV=Copy-number variants (global R^2^ estimates, derived from random vectors and from the Eigen-ANOVA method are shown in Table 3).

## 4 Discussion

Modern genomic data sets often combine information from multiple data-layers (e.g., pedigree, DNA-sequence, epigenomic information, gene expression, proteomics, metabolomics). The biological processes underlying these data can lead to complex dependencies between data-layers. MANOVA can be used to quantify the proportion of variance explained in multivariate settings. However, MANOVA is based on least-squares projections which are not-well suited for analysis of high-dimensional data. Reduced-rank regression methods (e.g. PLS, CCA) and penalized regressions (e.g., Lasso-type methods) can be used to confront the problems emerging when the number of features exceed sample size (p>>n). However, these methods are not adequate for estimation of proportion of variance explained, because they rely on regularization decisions (e.g., choosing the number of dimensions, or selecting a penalization parameter that controls sparsity) which are often based on cross-validation procedures that are not well-suited for inferences.

To overcome the limitations of existing methods, we developed two procedures to estimate the proportion of variance explained in settings where both the input and output sets are high-dimensional. The proposed approach uses random effects Gaussian models to estimate the proportion of variance of (independent) vectors in the linear span of an output set (***X***) that can be explained by regression on an input set (***W***). The resulting R^2^ estimate is a weighted average of the R^2^ values obtained for independent vectors. We considered two approaches to generate independent vectors in the linear span of the output set: The first one (MC-ANOVA) is a Monte Carlo method that uses randomly generated (independent) vectors in the linear span of the output set. The second one (Eigen-ANOVA) uses an orthogonal basis for the linear span of ***X***.

The two proposed methods have four important features: (i) Both methods can be used to perform analysis of variance when both explanatory and dependent data are high-dimensional; (ii) Estimates are entirely based on the likelihood function and there is no need to make regularization decisions (number of dimensions, penalty parameters). (iii) For any pair of information sets, the analysis of variance is not necessarily symmetric; therefore, the approach accommodates cases where the proportion of variance of ***W*** explained by ***X*** is not equal than the reciprocal. (iv) Finally, in addition to producing an R^2^ estimate, the proposed methods can shed light on important aspects of the underlying association patterns (e.g., decomposition of the global R^2^ on eigen-vector specific R^2^’s, distribution of R^2^ over possible vectors in the linear span of the output set).

Our simulations suggest that MC-ANOVA renders nearly unbiased estimates of the proportion of the variance of one set that can be explained by another. However, the Eigen-ANOVA exhibited small but systematic biases in scenarios in which the true proportion of variance was either too low or very high. The biases of Eigen-ANOVA were comparable, in magnitude, with those of the PLS regression. Therefore, for estimation of proportion of variance explained we recommend using MC-ANOVA.

The MC-ANOVA has clear computational advantages relative to Eigen-ANOVA and PLS because this method can render relatively accurate estimates of R^2^ based on a few hundreds of random vectors. Therefore, the number of regressions that one may need to perform can be much smaller than with PLS and the Eigen-ANOVA.

Consistent with our simulation results, the analyses of experimental data showed that in problems involving a high R^2^ the Eigen-ANOVA method yielded lower estimates of the proportion of variance explained than those obtained with the MC-ANOVA (e.g., see Figure 1 and Table 3). Inspection of the results of the Eigen-ANOVA for individual eigenvectors suggests that the downward bias of the method may originate from corner solutions (zero-estimates of R^2^) for eigenvectors associated with small eigenvalues. Therefore, if the only goal is to estimate the proportion of variance of one set explained by another set, we recommend using the MC-ANOVA method.

The Eigen-ANOVA method yields R^2^-values for each of the eigenvectors of the output set. This information can help elucidate whether global patterns (e.g., those associated with the top-eigenvectors) in one information set can be predicted from information contained in another information set. For instance, our analysis of the multi-omic breast cancer revealed that the patterns described in the top-eigenvectors derived from GE and ME are very similar; therefore, one should not expect big differences in tumor classifications that are based on the top-eigenvectors derived from either set. Interestingly, we found that in the analyses of omic data the R^2^ of individual eigenvectors showed a very sharp phase transition, suggesting that eigenvectors associated with intermediate and small eigenvalues may describe omic-specific patterns, or perhaps measurement error associated to each of the techniques.

The MC-ANOVA method can be used to characterize the distribution the R^2^ estimates across vectors in the linear span of the output set. We used this feature to study the effect of the trait-architecture; on the distribution of the R^2^ estimates. Our results indicate that while the average R^2^ does not depend on the distribution of the coefficients used to form random vectors (i.e., the *α*′*s*), the dispersion of the distribution is highly dependent on the process used to generate the weights. More specifically, random vectors that have non-zero weights on a small number of vectors have a larger dispersion in the distribution of the R^2^, compared to the dispersion observed when the random vectors have non-zero weights for all the basis in the linear span.

An important feature of the methods proposed in this study is that the R^2^ measure is not symmetric, in contrast to CCA. Our simulation study shows that if the underlying patterns are non-symmetric (e.g., when one of the linear spaces is a subspace of the other) the proposed estimation methods (in particular the MC-ANOVA) can detect the lack of symmetry very well (see Table 2). Interestingly, our analysis of multi-omic data from breast cancer patients exhibited cases where R^2^ was rather symmetric (e.g., the regression ME∼GE and the regression ME∼GE) and others that were highly asymmetric (e.g., CNV∼GE and GE∼CNV). The asymmetric cases suggest that almost all the variability in CNV can be predicted from GE (and ME as well); however, only a fraction of the GE variance can be explained by differences in CNV patters. This result is consistent with the hypothesis that most CNV have an impact on GE, but GE is also affected by factors other than CNV (e.g., methylation, environmental effects).

## In conclusion

We developed two methods for estimating the proportion of variance explained in problems in which both the input and output sets are high-dimensional. The MC-ANOVA method provided nearly unbiased estimates across a range of simulation scenarios. In addition to providing estimates of proportion of variance explained, the two methods can yield useful insight into the association patterns underlying multi-layered high-dimensional data.

