## Supplemental Figure for "Analysis of variance when both input and output sets are high-dimensional"

**I-Supplementary Figures**

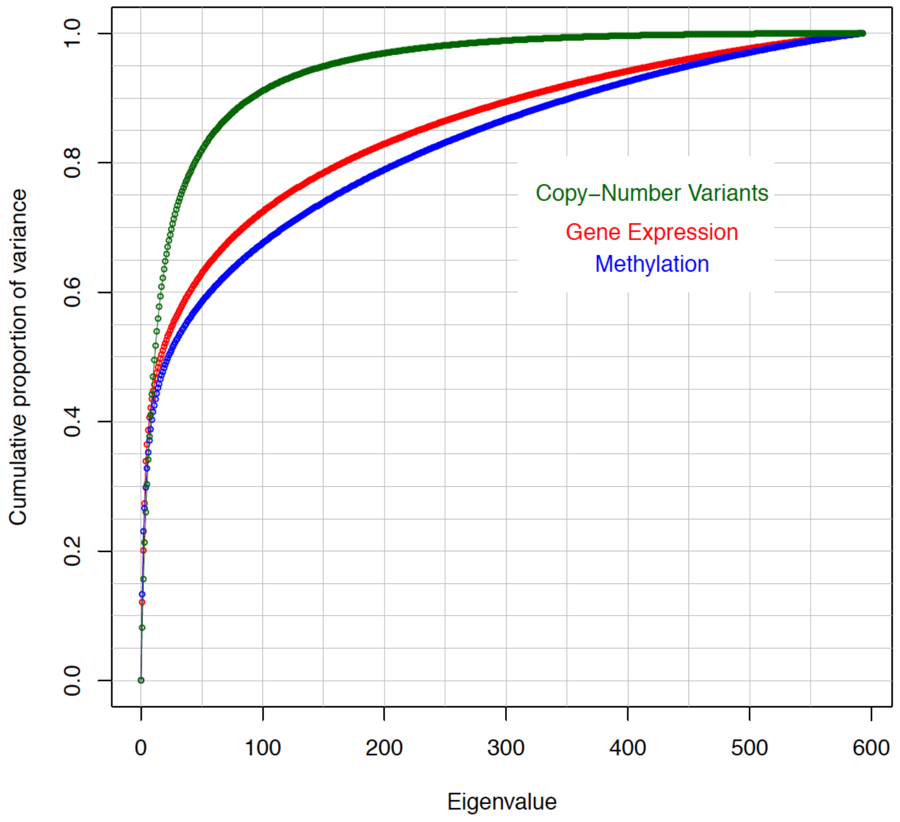


**Figure S1**: Cumulative proportion of variance explained by eigenvectors of each omic set. The cumulative proportion of variance was $\tau_j = \sum_{i=1}^{i=j} \lambda_i / \sum_{i=1}^{i=n} \lambda_i$ where $\lambda_i$ is the ith eigenvalue and $\sum_{i=1}^{i=n} \lambda_i$ is the sum of all the eigenvalues (j=1,…,n).
